# The Fc-mediated effector functions of a potent SARS-CoV-2 neutralizing antibody, SC31, isolated from an early convalescent COVID-19 patient, are essential for the optimal therapeutic efficacy of the antibody

**DOI:** 10.1101/2020.10.26.355107

**Authors:** Conrad E.Z. Chan, Shirley G.K. Seah, De Hoe Chye, Shane Massey, Maricela Torres, Angeline P.C. Lim, Steven K.K. Wong, Jacklyn J.Y. Neo, Pui San Wong, Jie Hui Lim, Gary S.L. Loh, Dong Ling Wang, Jerome D. Boyd-Kirkup, Siyu Guan, Dipti Thakkar, Guo Hui Teo, Kiren Purushotorman, Paul E. Hutchinson, Barnaby E. Young, David C. Lye, Jenny G. Low, Paul A. MacAry, Hannes Hentze, Venkateshan S. Prativadibhayankara, Kantharaj Ethirajulu, Damian O’Connell, Jason Comer, Chien-Te K. Tseng, Alan D.T. Barrett, Piers J. Ingram, Trevor Brasel, Brendon J. Hanson

**Author notes:** Corresponding author. (B.J.H.); (T.B.); (P.J.I.), (C.C.E.Z).

## Abstract

SARS-CoV-2-neutralizing antibodies are promising therapeutics for COVID-19. However, little is known about the mechanisms of action of these antibodies or their effective dosing windows. We report the discovery and development of SC31, a potent SARS-CoV-2 neutralizing IgG1 antibody, originally isolated from a convalescent patient at day 27 after the onset of symptoms. Neutralization occurs via a binding epitope that maps within the ACE2 interface of the SARS-CoV-2 Spike protein, conserved across all common circulating SARS-CoV-2 mutants. In SARS-CoV-2 infected K18-human ACE2 transgenic mice, SC31 demonstrated potent survival benefit by dramatically reducing viral load concomitant with attenuated pro-inflammatory responses linked to severe systemic disease, such as IL-6. Comparison with a Fc-null LALA variant of SC31 demonstrated that optimal therapeutic efficacy of SC31 requires intact Fc-mediated effector functions that can further induce an IFNγ-driven anti-viral immune response. Dose-dependent efficacy for SC31 was observed down to 5mg/kg when dosed before the activation of lung inflammatory responses. Importantly, despite FcγR binding, no evidence of antibody dependent enhancement was observed with the Fc-competent SC31 even at sub-therapeutic doses. Therapeutic efficacy was confirmed in SARS-CoV-2-infected hamsters, where SC31 again significantly reduced viral load, decreased lung lesions and inhibited progression to severe disease manifestations. This study underlines the potential for significant COVID-19 patient benefit for the SC31 antibody that justifies rapid advancement to the clinic, as well as highlighting the importance of appropriate mechanistic and functional studies during development.

**One Sentence Summary:** Anti-SARS-CoV-2 IgG1 antibody SC31 controls infection *in vivo* by blocking SP:ACE2 binding and triggering a Fc-mediated anti-viral response.

---

In December, 2019, a cluster of novel pneumonia (later named COVID-19) cases emerged and rapidly spread through human-to-human transmission in the city of Wuhan, China [1, 2]. High throughput sequencing of patient-derived samples revealed a novel beta-coronavirus, subsequently named SARS-CoV-2, as the etiological agent. SARS-CoV-2 was found to have 79.6% sequence homology to SARS-CoV, the virus responsible for an epidemic that caused 774 fatalities during 2002-2003 [3–6]. Like SARS-CoV, SARS-CoV-2 has the potential to cause severe respiratory distress and significant mortality and morbidity [1, 2]. While its natural reservoir remains unknown, based on sequence homology, SARS-CoV-2 is likely of bat origin [7]. SARS-CoV-2 was found to bind to angiotensin converting enzyme 2 (ACE2), the same cellular surface receptor used by SARS-CoV, via the receptor binding domain (RBD) of the viral surface Spike protein (SP) [8]. There is no pre-existing immunity to SARS-CoV-2 due to its low homology to circulating endemic coronaviruses. This, coupled with its high human-to-human transmissibility, has led to an on-going global pandemic that has currently caused more than 40 million infections worldwide and over one million fatalities.

Antibodies derived from the memory B cells of recovered patients have become an attractive approach to developing therapeutic antibodies for infectious disease. Such antibodies have previously been found to be protective against coronavirus diseases, such as SARS and MERS, in animal models [9, 10]. In particular, antibodies that blocked the viral SP protein from binding to ACE2 were highly potent at preventing infection [11, 12]. While such antibodies were not used clinically to treat coronavirus infections, antibodies derived from recovered patients had been successfully used to treat other infectious diseases, including the highly lethal Ebola virus disease with results superior to small molecule antivirals [13]. This indicates the highly promising therapeutic potential of antibodies derived from convalescent patients, especially those that inhibit viral entry through the functional ACE2 receptor. The SARS-CoV-2 SP protein and its RBD have also become the focus of numerous accelerated vaccine development programs [14–16]. However, given the challenges associated with large-scale roll-out of effective vaccines to the global population other options for protective immunity, even for shorter periods of time, must be considered. Therapeutic and prophylactic antibodies specific to SARS-CoV-2 have the potential to provide a viable treatment option before an effective vaccine is available, and further, can provide a much-needed treatment option for susceptible individuals who respond poorly to vaccination.

Multiple neutralizing antibodies against SARS-CoV-2 with therapeutic potential have been reported, with the most potent typically RBD-specific functioning by inhibition of binding to the ACE2 receptor [17–21]. Several have been tested in murine, hamster and non-human primate (NHP) models, where they and demonstrated both prophylactic and therapeutic efficacy and a smaller number are currently progressing through clinical trials [22–24]. In these pre-clinical models, therapeutic benefit was associated with reduced inflammation in the lung and correspondingly lower levels of proinflammatory cytokines and chemokines such as IL-6, CCL2 and CXCL10, which have been observed to be elevated during the COVID-19 cytokine storm [25]. Despite this, Antibody-Dependent Enhancement (ADE) of disease severity remains a major concern for the use of anti-viral antibodies as therapeutics. ADE can occur if Fcγ Receptor (FcγR) engagement mediates an increase in the infection of phagocytic cells that take up opsonized viral particles [26]. Indeed, ADE has been observed in both *in vitro* and *in vivo* studies of SARS infection [27–29]. Using mouse models, ADE has also been proposed to be a driver of the immune dysregulation observed in severe COVID-19 cases [26, 28], and therefore represents a risk for antibodies identified from a human immune response. Concerningly, in COVID-19 patients, higher anti-SP serum IgG levels have also been shown to correlate with hospitalization and severe disease [30, 31], however, there is no evidence to date that administration of convalescent plasma can lead to ADE.

As a result, several of the ongoing SARS-CoV-2 antibody programs have chosen to use Fc isotypes that do not engage FcγR, including the natural isotype IgG4 and engineered variants such as those carrying the LALA mutations [22]. However, this may be counterproductive as the signaling mechanisms underpinning the efficacy of these antibodies have not been evaluated, particularly the potential for beneficial engagement of FcγR to induce a targeted anti-viral response.

We present here the discovery and development of a potent RBD-binding neutralizing IgG1 antibody, SC31, originally isolated from an early convalescent patient. We have investigated the risk of ADE, and the impact of Fc functionality on its therapeutic efficacy via comparison with a FcγR null binding LALA variant and demonstrated that engagement of Fc receptors can drive a targeted anti-viral response, does not cause ADE and is required for complete efficacy of SC31 in rodent models of severe disease.

## Isolation and development of a potent neutralising antibody, SC31, from an early convalescent patient sample

Anti-SARS-CoV-2 antibodies were generated by single B cell antibody interrogation. SP-binding IgG^+^ B cells were sorted directly from patient PBMC samples obtained at 15- and 27-days post-symptom onset and cultured to induce antibody secretion. Despite sampling soon after the onset of symptoms, SP-specific IgG B cells were detected at a frequency below 1% of total IgG^+^ B cells (Fig. S1A). Analysis of antibody binding to wild-type SP ectodomain in the B cell culture supernatants identified a total of 36 SP binding antibodies of which ten had significant neutralizing activity (Fig. S1B). Heavy and light chain antibody pairs were isolated from these clones and converted to full IgG1 antibodies.

SC31, isolated from the patient at day 27 post-symptom onset, was the most potent neutraliser of SARS-CoV-2 infection of Vero E6 cells virus infection with an IC_50_ of 0.27μg/ml which equates to 1.85nM (Fig. S1C). Significantly, a comparison with IgG purified from the plasma fraction of the corresponding early convalescent patient sample showed that SC31 was 2000-fold more potent (Fig. 1A). SC31 still showed complete and strong neutralization of Vero E6 infection when the quantity of SARS-CoV-2 used in the neutralization assay was increased 10-fold to 1,000 TCID_50_ (Fig. 1B). SC31 bound to both the SP ectodomain and the RBD of SARS-CoV-2 with similar affinity (Fig. 1C), indicating that inhibition of receptor binding is likely the mechanism of neutralization. Indeed, SC31 demonstrated concentration-dependent inhibition of the interaction of both SARS-CoV-2 SP ectodomain and RBD with human ACE2 (Fig. 1D). Taken together, SC31 is a potent neutralizing antibody that acts by inhibiting the interaction of SP RBD with ACE2.

**Fig 1.**
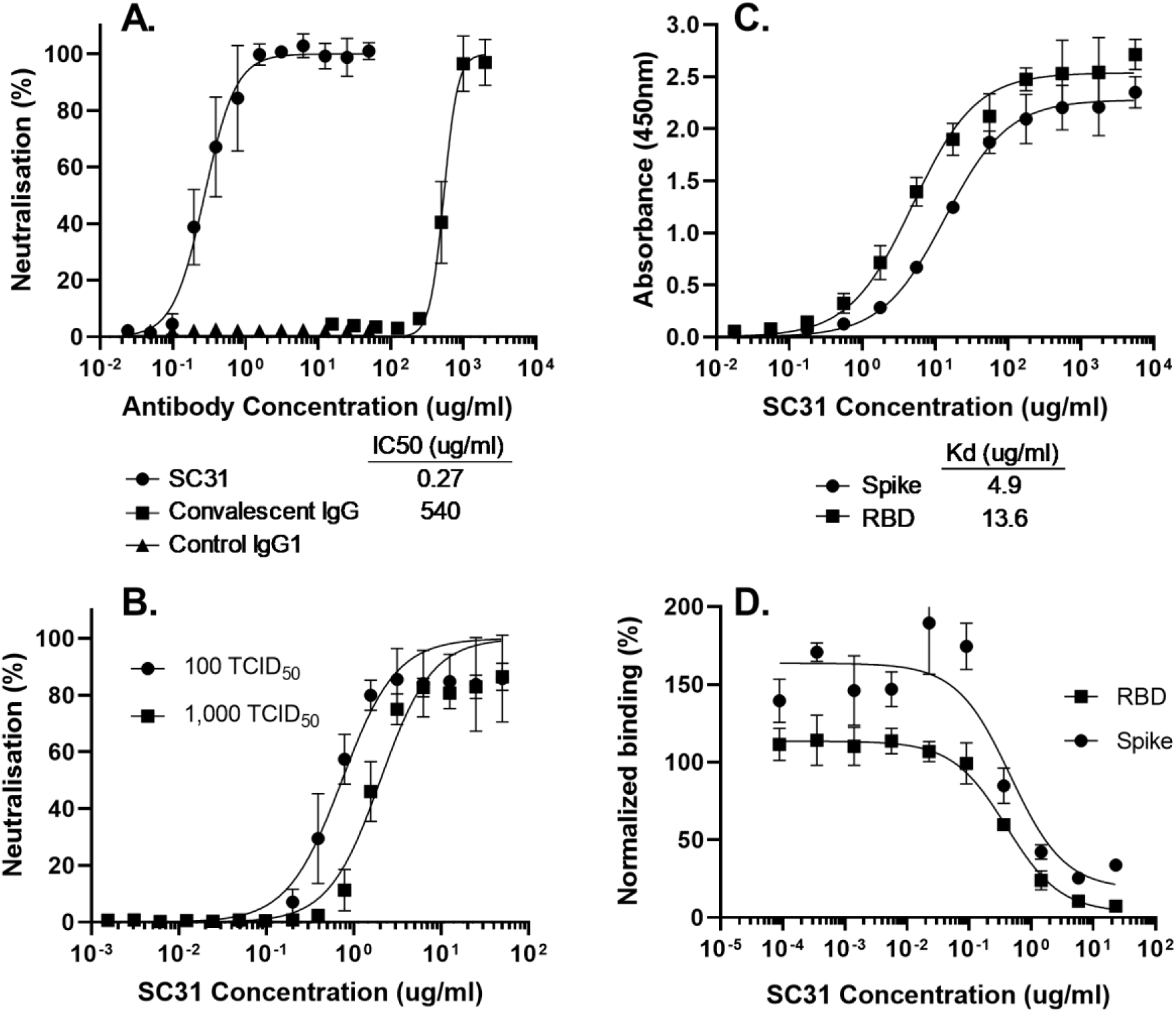
SC31 binds SARS-CoV-2 SP and neutralizes virus through inhibition of binding to ACE2. (**A**) Neutralization with 100 TCID_50_ of infectious virus by SC31 IgG1 in comparison to control IgG1 or IgG purified from donor serum. (**B**) Neutralization of 100 or 1000 TCID_50_ infectious virus by SC31 IgG1. Neutralization efficacy is represented as a percentage relative to uninfected and virus only controls. (**C**) Binding affinity of SC31 IgG to purified WT-spike or RBD as measured by ELISA. (**D**) Inhibition of WT-spike or RBD binding to cells expressing membrane-bound ACE2 by SC31 IgG1 at different concentrations as measured by flow cytometry. Binding is expressed as a percentage relative to no antibody or no viral protein controls. All results represent the mean of three independent replicates with bars showing standard error.

To accelerate the translation from the discovery of SC31 as a potential therapeutic to a clinical development candidate, rapid development activities were conducted. Developability optimization was performed to remove known sequence liabilities, resulting in a lead sequence with significantly improved charge variant profile (Table S1). Parallel establishment of high titer monoclonal master cell banks for multiple development variants of SC31 provided maximal optionality, pending functional and developability assessments, while producing sufficient material for *in vivo* functional studies, GLP toxicology studies, as well as for the process and analytical method development. A scale-up process for the lead master cell bank (MCB) was rapidly developed that robustly yielded a gross titer above 4 g/L at large scale GMP manufacture.

## The epitope of SC31 is located at the ACE2 interface of the SARS-CoV-2 SP protein and conserved in all circulating SARS-CoV-2 mutants

To investigate the structure and function of the epitope recognized by SC31, the binding of SC31 to naturally occurring SP mutants was evaluated. A total of 36 single amino acid SP mutations identified from publicly available databases were tested [32, 33]. We focused on 23 mutations that were within the receptor binding motif (RBM, aa438-506) and predicted to make direct contact with ACE2 receptor, or that were high frequency mutations beyond the RBM. The majority of these mutations were found to have minimal effect on SC31 affinity to SP and its binding to the ACE2 receptor (Fig. S2). Significantly, the two most common circulating mutations within the RBM, N439K and S477N, as well as the D614G mutation which delineates a major viral clade and is by far the most frequent SP mutation did not cause any significant changes in affinity (Fig. 2A). Six mutations (I434K, S438F, D467V, L455F, A475V, N501Y) were identified that resulted in significant loss of SC31 binding. Three of these six mutations, I434K, S438F and D467V, also resulted in significant or total loss of binding to the ACE2 receptor (Fig. 2B). This suggests that the loss of binding was due to destabilisation of the entire RBM structure and that these mutants would be unlikely to propagate further possibly due to reduced fitness. The other three mutations minimally reduced or even enhanced ACE2 receptor binding, as in the case of N501Y, and could be expected to lead to viral escape due to an alteration of the binding epitope. The locations of these three mutations are highlighted on a recently published crystal structure [34] and are likely to represent part of the SC31 binding epitope (Fig. 2C). Critically, none of these 3 SP mutants which appeared early on in the pandemic appear to be commonly circulating, perhaps due to other negative effects on the stability of the protein. SC31 therefore binds to a stable epitope in the RBD of the SARS-CoV-2 SP protein that is well conserved in all commonly circulating SARS-CoV-2 variants.

**Fig. 2.**
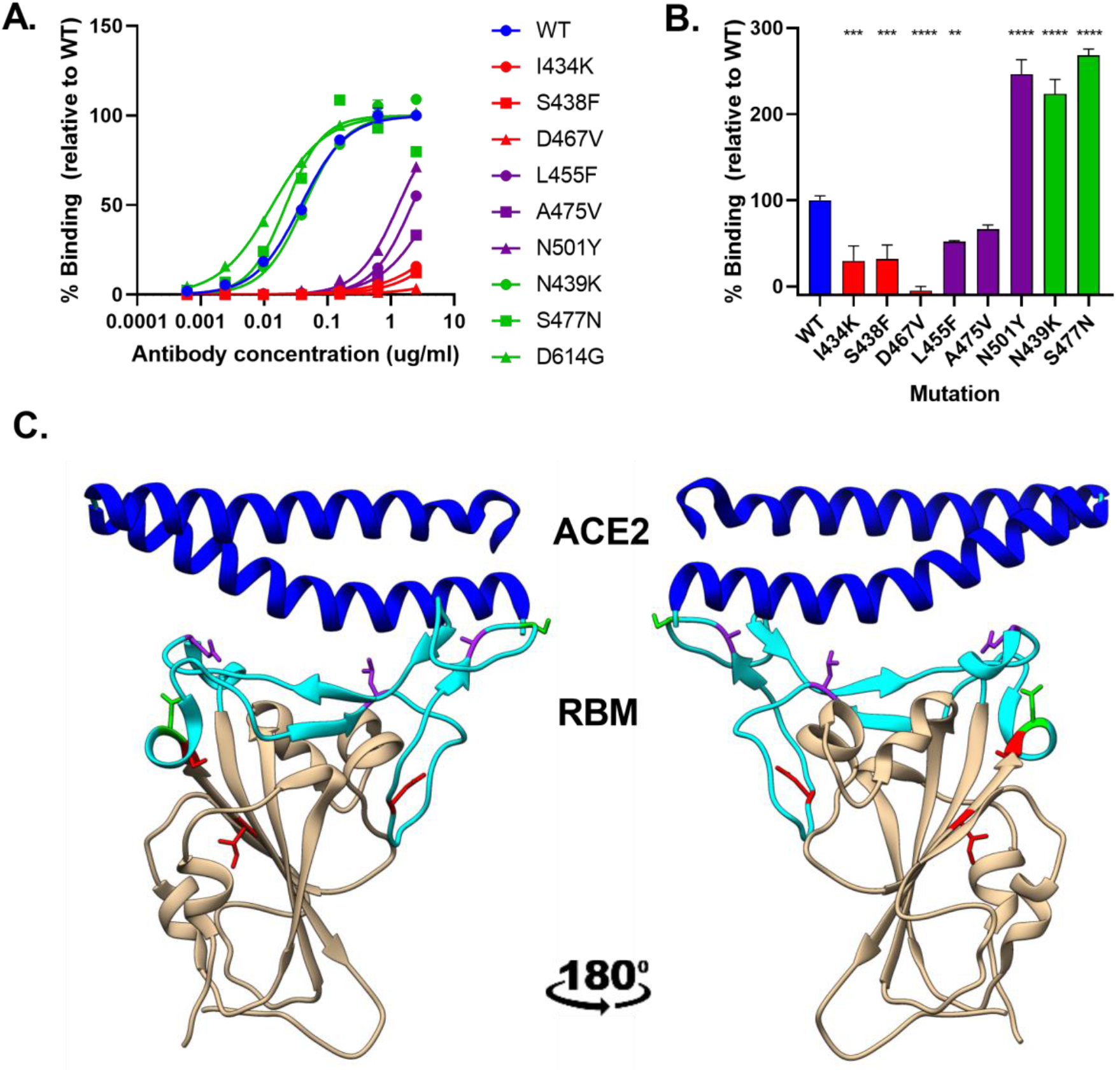
SC31 binding to SP variants identifies its ACE2 interface epitope. (**A**) Binding affinity of SC31 as determined by ELISA to purified wild-type spike (blue) and spike mutants that either do not affect SC31 or ACE2 binding (green), affect SC31 but not ACE2 binding (yellow) or affect both SC31 and ACE2 binding (red). Results are the mean of three independent replicates and are represented as a percentage of maximal absorbance against wild-type spike at the highest antibody concentration. (**B**). Binding affinity of purified wild-type and mutant spike protein to hACE2-expressing CHO cells based on fluorescence intensity measured by flow cytometry. Results are the mean of three independent replicates with bars showing the standard error and are represented relative to wild-type spike binding to ACE2. Statistical significance was determined vs. wild-type for each mutant by one-way ANOVA: ** p<0.01, *** p<0.001, **** p<0.0001. (**C**). Location of single amino acid mutations (green/yellow/red) on a crystal structure of RBD showing the interaction of the RBM (cyan) with ACE2 N-terminal helix (blue).

## SC31 requires Fc-mediated effector mechanisms for maximal therapeutic efficacy in the SARS-CoV-2 K18-hACE2 mouse severe disease infection model but does not cause ADE

SC31 is an IgG1 antibody therefore binding of the Fc domain to FcγR on immune cells is expected. FcγR binding may stimulate beneficial Fc-mediated effector function, including Antibody-dependent Cellular Cytoxicity (ADCC), Antibody-dependent Cell Phagocytosis (ADCP), Antibody-dependent cell-mediated virus inhibition (ADCVI) and Complement-Dependent Cytoxicity (CDC), but, as previously discussed, may also lead to ADE for viral infection. ADE is especially important when antibodies reach sub-neutralising concentrations. To ameliorate the risk of ADE for anti-viral antibodies, it is possible to utilize an Fc isotype that does not bind to FcγRs, however, this could also impact the potency of the antibody if Fc-mediated effector functions contribute to the therapeutic efficacy of the antibody.

To first establish if Fc-mediated effector function could have a role in the therapeutic efficacy of SC31, the ability to induce Fc effector-mediated activity was investigated by comparing against an FcγR null-binding variant (LALA) of the antibody. The upstream activation of the FcγRIIIa ADCC signalling pathway was evaluated using a Jurkat reporter cell line co-cultured with target HEK293 cells expressing membrane-bound SARS-CoV-2 SP. Here, SC31 was confirmed to induce dose-dependent activation of ADCC signalling. In contrast, there was no activation of ADCC observed with the FcγR null-binding variant (LALA) of the antibody, similar to the mock-transfected HEK293 cells (without SP) (Fig. 3A and B) [35].

**Fig 3.**
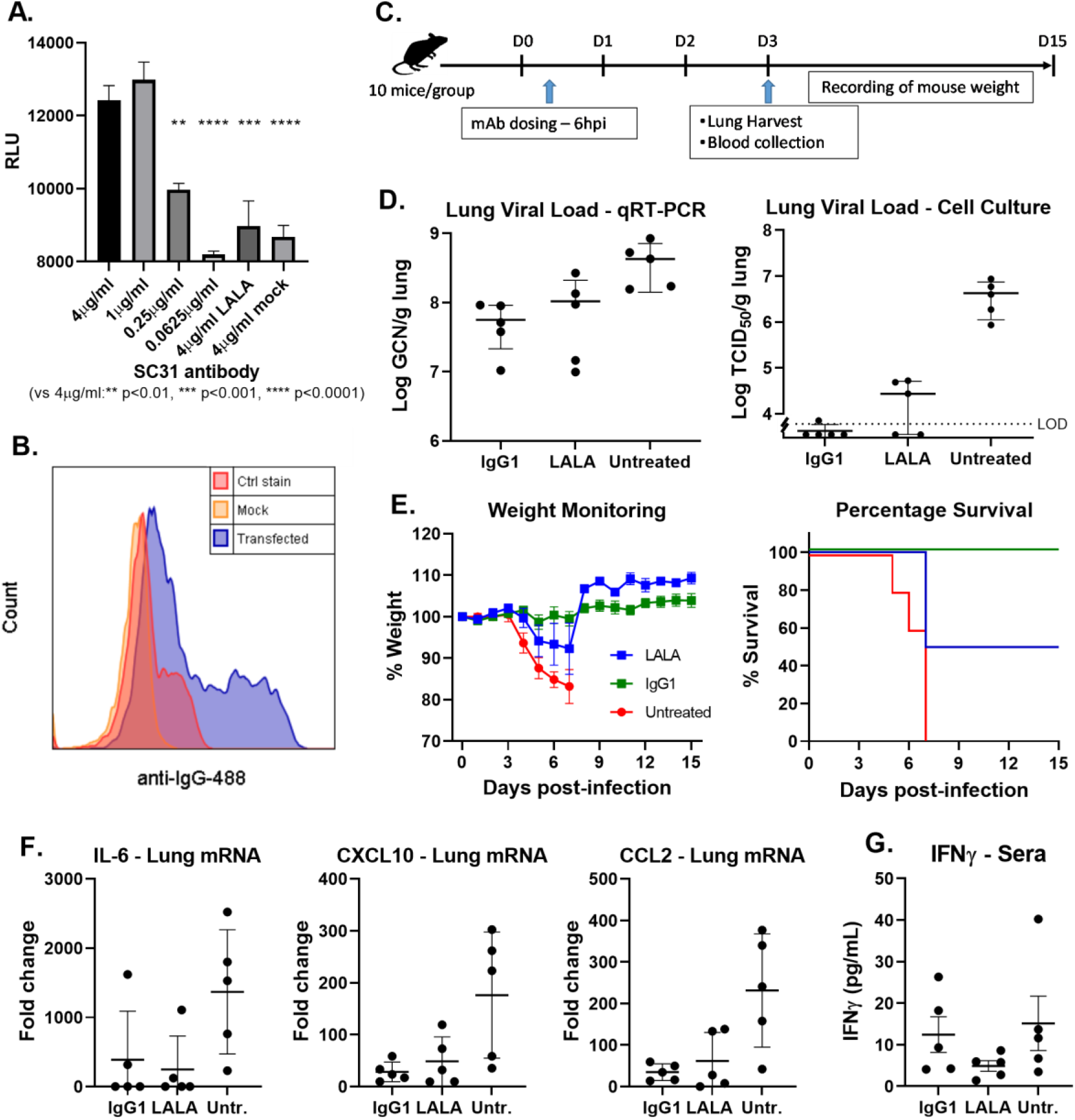
SC31 requires Fc effector functions for optimal therapeutic benefit. (**A**) Activation of ADCC signaling by SC31 or null-binding LALA variant at various concentrations incubated with FcγRIIIa-expressing reporter cell line and spike or mock transfected HEK293, as determined by luciferase expression. Statistical significance in comparison to the highest SC31 concentration of 4ug/ml was determined by one-way ANOVA. (**B**). Specific binding of SC31 to spike transfected HEK293 cells in comparison to mock transfected cells or fluorophore stain only (Ctrl). (**C**) Overview of therapeutic study design with SC31 IgG1 and LALA variant antibodies. Antibodies were dosed at 20mg/kg 6hpi and 5 mice sacrificed at 3 dpi to ascertain lung viral and cytokine load as well as antibody serum titers with the remainder monitored for weight and survival. (**D**) Lung viral load of IgG1-, LALA-treated or untreated mice at 3 dpi as measured by genome copies (left) or infectious virus (right). The limit of detection (LOD) is indicated by the dotted line. (**E**) Disease progression in infected mice as indicated by weight loss (left) or survival (right). (**F**) Lung cytokine mRNA expression determined by qRT-PCR and represented as fold-change over uninfected mice. (**G**) Sera IFNγ protein level determined by sandwich ELISA. Each point represents one individual mouse and all error bars show standard error.

Subsequently, the therapeutic efficacy of SC31 and its LALA variant were evaluated in a SARS-CoV-2 K18-hACE2 mouse model to investigate if Fc effector mechanisms contributed to therapeutic efficacy. Severe disease manifestations with SARS-CoV-2 infection have been demonstrated in K18-hACE2 transgenic mice [21, 36–38]. Mice infected intranasally with 1.2 × 10^4^ TCID50 (nCoV-19/Singapore/3/2020) presented with severe disease including lethargy, weight loss, overexpression of proinflammatory cytokines/chemokines (IL-6, CXCL10, and CCL2) and, ultimately, death between 6- and 8-days post infection (dpi) with associated high virus titers in lung tissue (Fig. S3).

SC31 and the LALA variant (20mg/kg) were introduced by intraperitoneal (I.P.) administration 6 hours post virus infection (hpi). At 3 dpi, lungs from half of the mice per group were harvested for total and infectious viral load quantification and cytokine/chemokine mRNA and protein expression analysis. The remaining mice were monitored for survival and weight for a further 15 days (Fig. 3C). Quantification of viral RNA and infectious virus showed that the LALA variant was somewhat less efficacious than the wild-type IgG1 in controlling viral load (Fig. 3D). However, in contrast, significantly more severe illness was observed in the virus-infected mice treated with the LALA variant, demonstrated by their greater weight loss and higher mortality (Fig. 3E). Observations of the pro-inflammatory cytokine IL-6 and chemokines CXCL10 and CCL2 from virus-infected mice showed that, while there was a similar reduction in IL-6 for both IgG1 and the LALA variant, the LALA mice showed significantly higher levels of CXCL10 and CCL2 to those treated with IgG1 (Fig. 3F). Where both data were available the protein levels mirrored the mRNA levels. Intriguingly the protein levels of IFNγ were markedly increased in the IgG1 group, suggesting that although the systemic pro-inflammatory response is decreased, there is likely a beneficial targeted anti-viral response. Taken together these results indicate that in addition to the ability to inhibit the binding of SARS-CoV-2 to ACE2, Fc-mediated modulation of the immune response is critical for optimal therapeutic benefit.

Finally, to investigate the risk of ADE, SC31 and its LALA variant were tested at sub-neutralizing concentrations using a SARS-CoV-2 pseudovirus and FcγRIIIa-expressing THP-1 and Raji cell lines. ADE has been previously been modelled for SARS-CoV pseudovirus in these same cell lines [28, 39]. Importantly, no pseudovirus infection was observed in both THP-1 and Raji cells for either antibody at all concentrations tested, indicating that SC31, despite its FcγR binding leading to potent Fc mediated effector functions, is unlikely to mediate ADE (Fig. 4A and B). It has been suggested that pH selective binding of anti-viral antibodies, i.e. lower binding affinity to the viral target at lower pHs may predict a risk of ADE as antibodies could dissociate from the virus in the low pH environment of the endosome and release the virus to enter the cell. We found that SC31 maintains a high affinity for SP down to pH 4.5 and this may explain the lack of observable ADE for this antibody (Fig 4C).

**Fig. 4.**
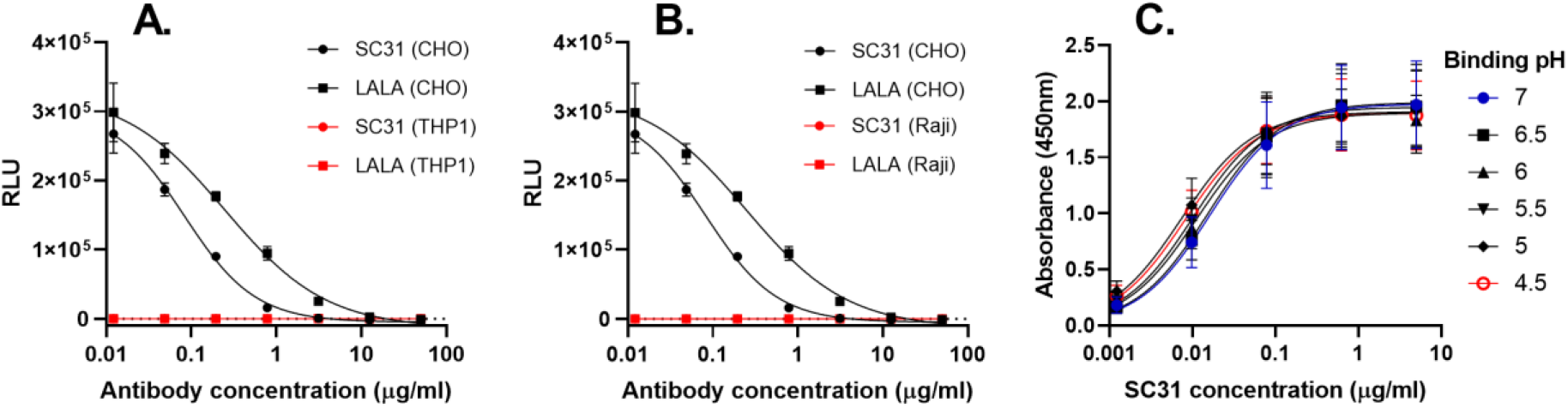
SC31 does not induce ADE. (**A**and **B**) Lack of SARS-CoV-2 pseudovirus infection co-incubated with SC31 or LALA variant in FcγRIIIa expressing cell lines THP-1 (A) and Raji (B) in comparison to ACE2-expressing CHO cell line based as determined by luciferase reporter gene expression. (**C**) Retention of SC31 binding affinity for WT-spike between pH4.5-7.0 as measured by ELISA. All results represent the mean of three or four independent replicates with error bars showing standard error.

## SC31 provides potent dose-dependent therapeutic benefit in the SARS-CoV-2 K18-hACE2 mouse severe disease infection model

Understanding the minimum therapeutic dosing level and optimal timing of an anti-SARS-CoV-2 antibody therapy for the treatment of COVID-19 remains critical to evaluate the potential of such an antibody in a clinical setting. To further evaluate the therapeutic potency and dose response of SC31, mice were treated with 2, 5, 10, or 20mg/kg doses 6 hpi. At 3 dpi, half the mice were sacrificed for quantification of lung viral RNA, infectious virus, and cytokine/chemokine expression. The remaining mice were monitored for survival and weight for a further 25 days (Fig. 5A). A dose-dependent reduction in weight loss, starting at 3dpi, was observed in all virus infected mice. A similar dose-dependent effect on survival was observed, with no mortality in animals treated with the highest dose (20mg/kg), 50% mortality in mice dosed with either 5 or 10mg/kg and no survival at the lowest dose (2mg/kg). All surviving mice overcame clinical signs of disease by day 13, as evidenced by the return to their pre-infection weight (Fig. 5B). Quantification of SARS-CoV-2 RNA in lung tissue showed a dose-dependent reduction of viral RNA ranging from 0.8 to 1.5-log from the lowest to highest dose, respectively (Fig. 5C). A corresponding reduction in infectious virus was observed with a greater than three log reduction with 5mg/kg antibody and down to the limit of detection for the majority of mice dosed at 20mg/kg (Fig 5D). In the 10 and 20mg/kg treatment groups, pro-inflammatory cytokine IL-6 and chemokines CXCL10 and CCL2 showed a decrease to levels similar to that in uninfected mice (Fig. 3E). Taken together, these results suggest that SC31 has therapeutic benefit at doses above 5mg/kg.

**Fig 5.**
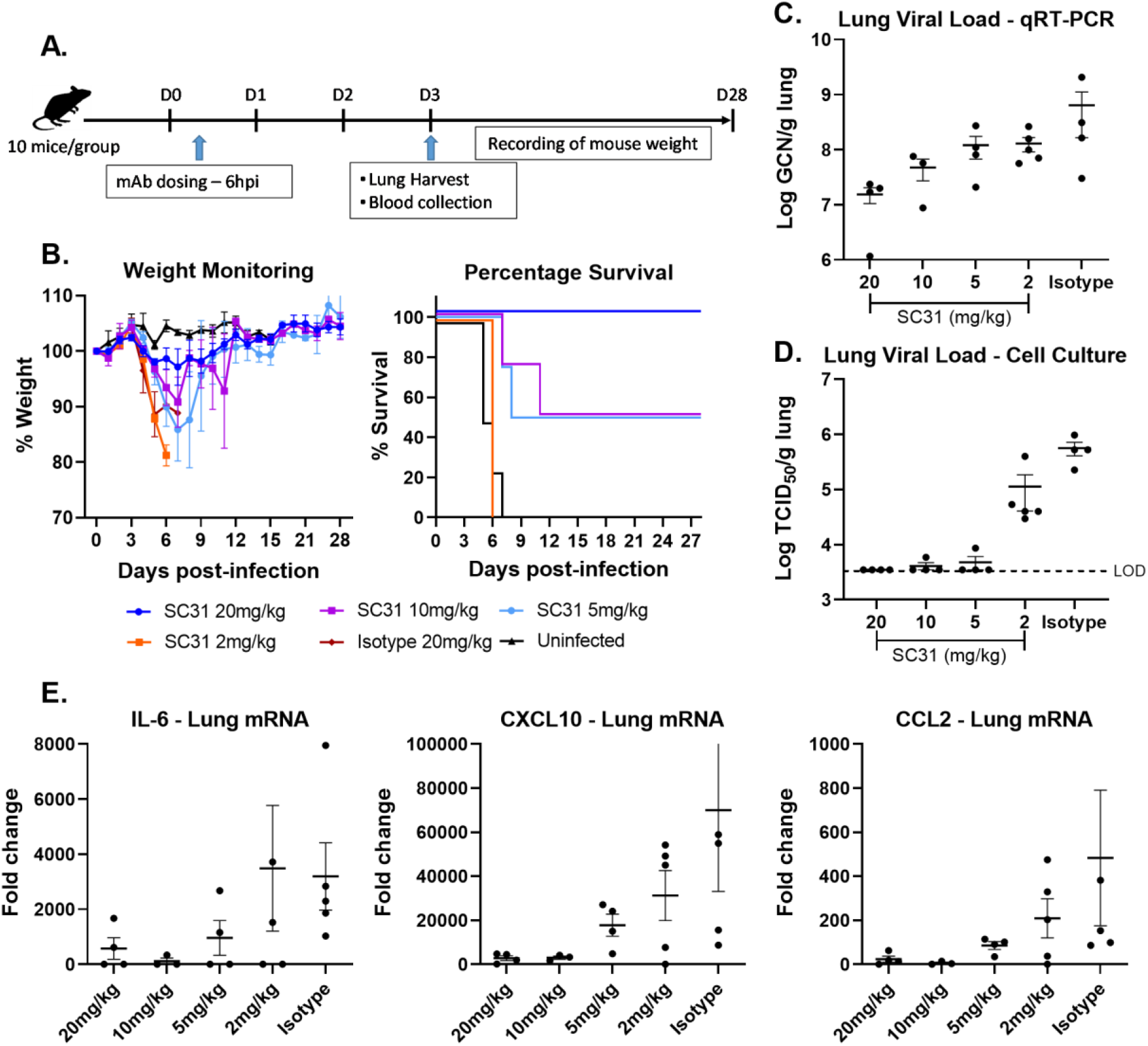
Dose dependent therapeutic benefit of SC31. (**A**) Overview of therapeutic study design with different SC31 IgG1 doses or isotype control at 20mg/kg. Antibodies were dosed at 6 hpi and 5 mice sacrificed at 3 dpi to ascertain lung viral and cytokine load as well as antibody serum titers with the remainder monitored for weight and survival. (**B**) Disease progression in infected mice as indicated by weight loss (left) or survival (right). (**C-D**) Lung viral load of antibody treated mice at 3dpi as measured by genome copies (C) or infectious virus (D). The limit of detection (LOD) is indicated by the dotted line. (**E**) Lung cytokine mRNA expression determined by qRT-PCR and represented as fold-change over uninfected mice. Each point represents one individual mouse and all error bars show standard error.

To ascertain the efficacious dosing window for SC31, mice were treated with 20mg/kg of antibody at 6, 24 and 48 hpi with half the mice sacrificed for lung viral load and cytokines at 3 dpi and the remainder monitored until Day 15 (Fig. 6A). All mice treated at 6 and 24 hpi survived with minimal weight loss while mice treated at 48 hpi lost weight and succumbed to disease at the same rate as untreated mice (Fig. 6B). Lung viral RNA, infectious virus and IL-6, CXCL10 and CCL2 levels followed the same trend with similar therapeutic benefit observed in mice treated at 6 and 24 hpi while mice treated at 48 hpi exhibited values similar to untreated mice (Fig. 6C-E). Taken together, the results indicate that the efficacious dosing window in this model is before 48 hpi, *i.e.*, prior to the peak of viral infection and inflammation that was also observed at day 3 (Fig. S3), suggesting that early clinical dosing before the onset on lung inflammation may be necessary for optimal therapeutic benefit.

**Fig 6.**
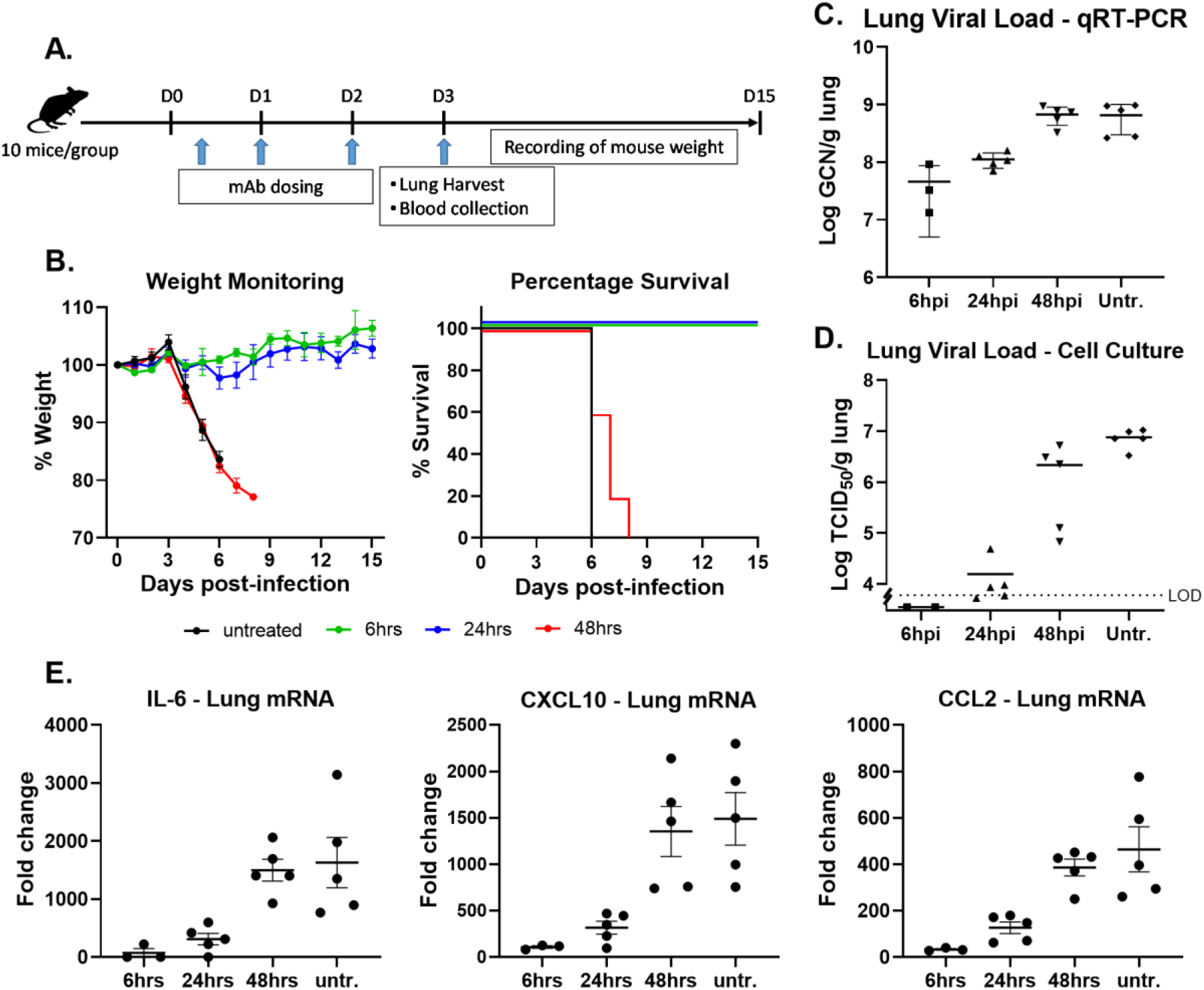
Efficacious dose window for SC31is before viral peak. (**A**) Overview of therapeutic study design with SC31 IgG1 doses at 20mg/kg at different timepoints. 5 mice were sacrificed at 3dpi to ascertain lung viral and cytokine load as well as antibody serum titers with the remainder monitored for weight and survival. (**B**) Disease progression in infected mice as indicated by weight loss (left) or survival (right). Lung viral load of antibody treated mice at 3 dpi as measured by genome copies (**C**) or infectious virus (**D**). The limit of detection (LOD) is indicated by the dotted line. (**F**) Lung cytokine mRNA expression determined by qRT-PCR and represented as fold-change over uninfected mice. Each point represents one individual mouse and all error bars show standard error.

## SC31 provides potent therapeutic benefit, protecting against severe COVID-19-like disease in Golden Syrian hamsters infected with SARS-CoV-2

To confirm the efficacy of SC31 in a second disease model, against a second SARS-CoV-2 virus strain, and investigate the ability of SC31 to inhibit progression to severe lung inflammation, the Golden Syrian hamster model was used. This model shows severe disease manifestations with SARS-CoV-2 infection, including predictable weight loss, alveolar inflammation and bronchiolo-alveolar hyperplasia [40, 41]; and has been used for the evaluation of other COVID-19 antibody treatment candidates [18, 42].

To evaluate the therapeutic efficacy of SC31, hamsters (6 male/6 female) were intranasally challenged with 5 × 10^5^ TCID_50_ SARS-CoV-2 (USA_WA1/2020) and half were treated with SC31 (20mg/kg). Animals were monitored daily, including body weight, subcutaneous body temperature and collection of nasal swabs for SARS-CoV-2 RNA and infectious virus analysis before lung harvest at 7 dpi. SC31 treated hamsters showed a transient rise in body temperature (day 1) and minor drop in weight that returned to pre-infection weight by day 6 (Fig. 7B and A). In contrast, vehicle control treated hamsters showed only a minor rise in temperature and continually lost weight through day 6, losing, on average, 10 percent of their pre-infection weight. When analyzed for both infectious and total viral load, nasal swabs showed a 1-2 log reduction, respectively (Fig. 7D and C). Lungs collected from the vehicle control treated hamsters 6-7 dpi consistently showed higher total viral load than those treated with SC31 (Fig. 7E) and presented with mottled, dark red lungs and 1-3 mm lesions diffusely distributed throughout the tissue, correlating histologically to hemorrhage and bronchiolo-alveolar hyperplasia, respectively (Fig. 7F and G). In stark contrast, all except one of the lungs collected from the treated group, did not show this gross phenotype.

**Fig 7.**
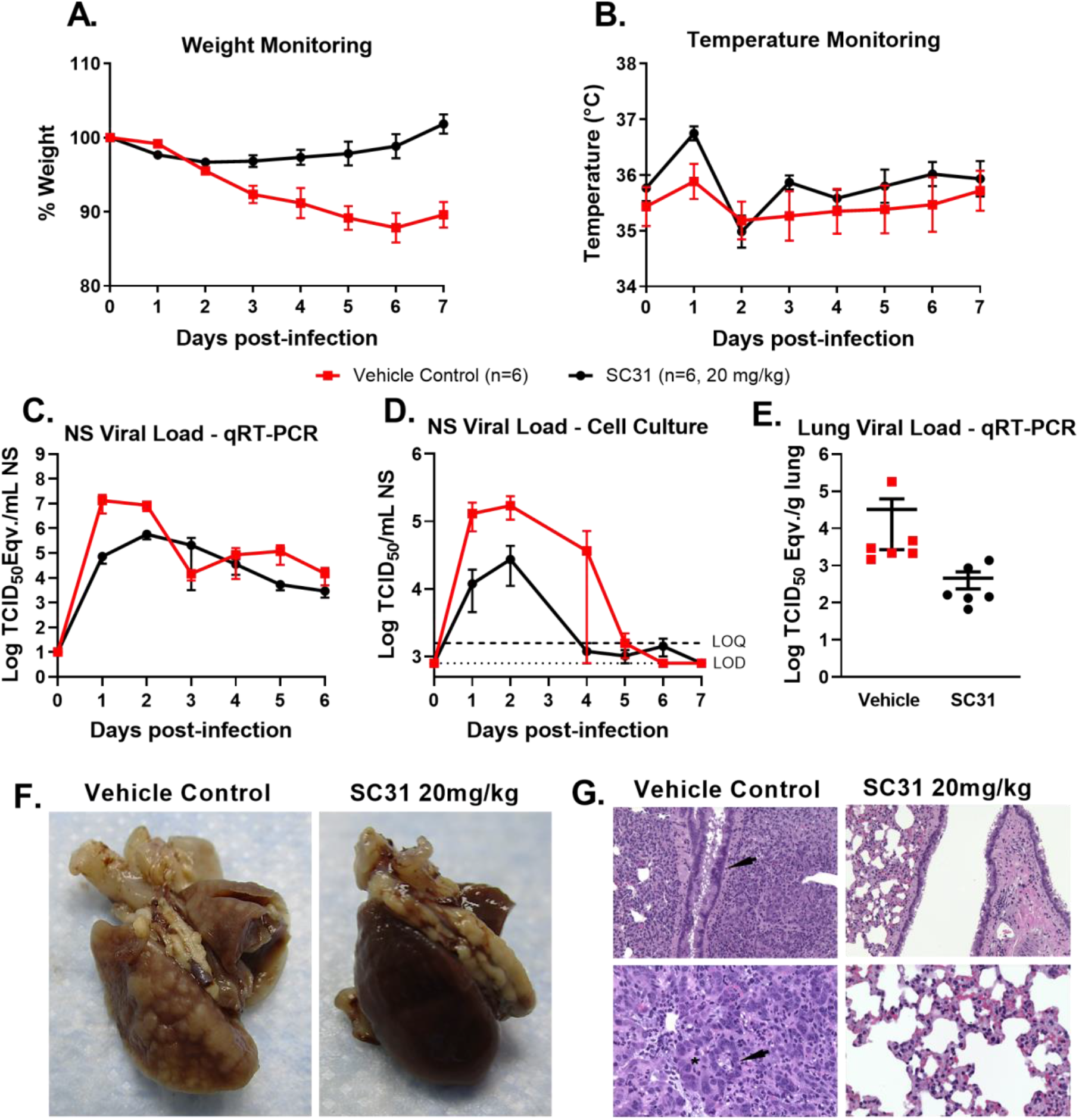
SC31 minimizes disease after SARS-CoV-2 challenge in Golden Syrian hamsters. Twelve hamsters (6 male/6 female) were intranasally challenged with 5 × 10^5^ TCID_50_ SARS-CoV-2; 6 animals were treated with SC31 antibody or vehicle as indicated at 4 hpi. (**A**) Percent body weight changes from baseline following SARS-CoV-2 intranasal challenge and SC31 treatment at 4 hpi. (**B**) Subcutaneous body temperatures as measured *via* implanted IPTT-300 temperature transponders. (**C**) SARS-CoV-2 RNA was lower in nasal swab (NS) samples collected from antibody-treated hamsters. (**D**) Infectious viral loads in NS as determined via TCID_50_ assay were lower in hamsters treated with SC31. (**E**) Viral RNA was 1-3 log lower in SC31-treated hamsters; infectious virus was not detected in lung samples collected 7 dpi. (**F)**Harvested lungs from vehicle control-treated (left) and SC31-treated (right) hamsters at 7 dpi. Images taken following removal of the left lobe for viral load analysis; representative images are shown (vehicle control female hamster 1, left image; SC31-treated male hamster 12, right image). (**G**) Histopathology examination of fixed lung tissue; representative images are shown (vehicle control female hamster 1, left images; SC31-treated male hamster 12, right images). For all panels, error bars show standard error.

Additional histopathological findings included an increase of type II pneumocytes indicative of acute lung damage, hyperplastic and hypertrophied bronchial/bronchiole epithelium (Figure 7G, top left, arrow) and alveoli lined by type II pneumocytes (top right, arrow) and enlarged nuclei (top right, asterisk). Conversely, microscopic findings associated with bronchial/bronchiole epithelium (Figure 7G, bottom left) and alveolar epithelium (Figure 7G, bottom right) were less severe or normal in SC31-treated hamsters. Results from this study demonstrate that SC31 can effectively treat acute viral infection and limit the development of severe lung inflammation in the Golden Syrian hamster model.

## Discussion

There is an urgent need for development of safe and efficacious therapeutic antibodies to address the global COVID-19 pandemic. Such antibodies represent a potentially critical treatment option in the absence of a widely available vaccine especially for elderly and other at-risk groups. Nevertheless, relatively few antibody candidates have progressed to clinical evaluation to date and only a few have reported strong efficacy data in pre-clinical models [18, 21, 24, 43]. This is, in part due to the challenges of demonstrating that any clinical candidate is both safe and potent enough to predict clinical benefit, which requires the rapid development and scale-up manufacture of a stable drug-like antibody, and evaluation in complex animal models. Further, concern over the possibility of ADE has led some efforts to focus on potentially sub-optimal Fc-effector silent candidates (IgG4 or antibodies with Fc mutations that abolish FcyR binding). Although data is still emerging, two IgG1 antibody programs [44, 45] are already reporting promising early data in clinical trials, but there is limited information on their mechanism of action and the possibility of ADE at sub-therapeutic doses is still an open question.

In this study we present the generation and characterization of a highly efficacious anti-SARS-CoV-2 IgG1 neutralizing antibody, SC31. The binding site of SC31 has been mapped to the receptor binding motif of the SP protein and has been shown to be conserved across all common circulating SARS-CoV-2 mutants, including the most common variant, D614G. In parallel to the rapid development and scale-up manufacture of the SC31 antibody, the role of Fc-mediated effector activity on the therapeutic efficacy of the antibody was evaluated alongside the potential to cause ADE in order to establish the optimal antibody format for evaluation in the clinical setting.

Although IgG1 isotype antibodies are predicted to bind stimulatory FcγR and trigger signaling as well as Fc-mediated effector functions, such as ADCC, ADCVI and CDC, not all IgG1 anti-SP antibodies exhibited the same capacity to elicit Fc-mediated effector functions. Specifically, it has been shown that some anti-SP IgG1 antibodies exhibited weak or even negligible ADCC activity [17, 46]. This may explain the variation between *in vitro* and *in vivo* potency reported for different anti-SP antibodies as well as a potential reason for the lack of immunity afforded by higher anti-SP-IgG titers in some COVID-19 patients [30]. This also suggests that the ability of an antibody to promote Fc-mediated effector functions and signaling may be another key factor in therapeutic efficacy in addition to neutralization potency. Importantly, the SC31 antibody was clearly able to trigger Fc-mediated effector functions, as evidenced by activation of ADCC signaling in contrast to a Fc-effector null LALA variant, but notably, despite this, SC31 showed no evidence of ADE at sub-therapeutic doses, identical to the LALA variant.

This ability of SC31 to induce Fc-mediated effector signaling was subsequently demonstrated to be essential for the optimal therapeutic efficacy. When SC31 and its LALA variant were evaluated in an *in vivo* mouse model of SARS-CoV-2 infection, the LALA variant was observed to have much reduced therapeutic efficacy with only 50% survival compared with 100% survival for SC31. This correlated with higher lung viral load as determined by both viral RNA and infectious virus, suggesting that the LALA variant was not able to clear virus, thereby triggering higher inflammatory responses. This indicates that FcγR engagement of lung phagocytic cells by SC31 not only did not result in excessive pro-inflammatory signaling, but appeared to induce a more targeted and robust anti-viral response characterized by higher IFNγ levels. The reduction in systemic pro-inflammatory markers including IL-6, CCL2 and CXCL10 is likely due to the reduction in viral load that dampens the systemic inflammatory signaling by various innate viral pathogen recognition receptors such as Toll-Like Receptor 7 [47]. The increase in IFNγ levels, however, may reflect the engagement of a targeted NK and T-cell driven anti-viral response, as these are the primary producers of IFNγ. These benefits were also observed in the hamster model where treatment with IgG1 resulted in significantly reduced lung pathology and a higher peak temperature compared the untreated group characteristic of a more robust anti-viral response. In summary, although a recent study showed that anti-SP IgG from COVID-19 patients with severe disease can promote hyperinflammatory responses in alveolar macrophages [48], our data indicates that SC31 is unlikely to pose a risk of systemic immune dysregulation when used as a therapeutic.

Severe disease and mortality in COVID-19 appears to be driven by excessive inflammation due to failure of the immune system to control viral infection in the lung [49]. SC31 showed potent dose-dependent therapeutic efficacy above 5mg/kg, however, once the inflammatory cascade is triggered, SC31 was no longer able to exert a therapeutic effect as evidenced by the poor outcome when mice were treated 48 hpi. This indicates that SC31 antibody therapy is best administered prior to the onset of severe symptoms.

Overall, we have demonstrated that the anti-SP IgG1 antibody SC31, generated from an early convalescent patient at day 27 after symptom onset, is able to control infection in two animal models of COVID-19 disease by decreasing viral load and protecting against lung damage, and has the potential to be a highly efficacious therapeutic in the clinical setting. This efficacy is driven by the dual mechanisms of potent neutralization of SARS-CoV-2 infection through blocking SP binding to the human ACE2 receptor, and induction of a robust anti-viral response driven by Fc-mediated effector functions, but importantly, without concomitant ADE. Furthermore, we have shown that SC31 is efficacious against two circulating strains of SARS-CoV-2. Following a highly accelerated non-clinical and CMC development program that has resulted in an efficient scaled-up manufacturing process yielding over 4g/L in large scale GMP manufacture, SC31, also known as HMBD-115 and AOD01 for development, will shortly begin human trials in COVID-19 patients.

## Supporting information

Chan et al Supplemental Information

## Acknowledgments

We gratefully acknowledge the assistance of Aw Lay Tin, DSO National Laboratories, for technical assistance with FACS and BSL3 respectively, and we thank Vithya Manoharan for assistance with data analysis and graphing. We are grateful for funding from Ministry of Defence, Singapore.

## Supplementary Materials

Materials and Methods

Figures S1-S3

Tables S1

