## Supplementary material for "The Fc-mediated effector functions of a potent SARS-CoV-2 neutralizing antibody, SC31, isolated from an early convalescent COVID-19 patient, are essential for the optimal therapeutic efficacy of the antibody": Chan et al Supplemental Information

### Supplementary Information

#### Materials and Methods

Single cell sorting and culture

Ficoll-Paque patient PBMCs at $5x10^6$cells/ml concentration were incubated with 10ug/ml Twin-strep-tagged WT-spike for 1hr at 4°C in FACS buffer (1x PBS, 5mM EDTA, 1% fetal calf serum), washed in 1xPBS and then stained with fluorescently labelled antibodies at the following concentrations (5µl anti-HuCD19-Pacific Blue, 5µl anti-HuCD27-Alexa647, 2.5µl-anti-HuIgG-BV711, 5µl anti-HuCD38-PE-Cy7, 2.5µl anti-Strep-tagII-Alexa488 in 100µl of FACS buffer per $10^6$ PBMC cells) for 30min at 4°C. Cells were washed with 1xPBS and resuspended in FACS buffer containing 1:100 propidium iodide (PI) before sorting on a FACSAria Fusion. $PI^-$, $CD19^+$, $IgG^{+,}$ $Spike^+$ cells were sorted individually into 96-well plates containing 50µl of IMDM culture medium, $5x10^3$ 3T3-msCD40L feeder cells [1] and supplements and cytokines at concentrations previously described without CpG and addition of IL-21 at 10ng/ml [2]. Cells were spun down after 8 days of culture and stored with 10µl of lysis buffer (QuickExtract RNA Extraction Solution, Lucigen) and supernatants tested by ELISA and microneutralization.

Antibody and viral protein expression and purification

Antibody variable heavy and light chain sequences were cloned into a pCMV-promoter driven expression plasmid containing the appropriate human IgG1 heavy, kappa or lambda light chain constant regions as well as a leader sequence for secretory expression. For spike protein expression, the complete ectodomain including the leader sequence (Accession No. MN908947, S gene amino acids 1-1208) together with a C-terminal Twin-strep tag (WSHPQFEK-GGGSGGGSGGS-SAWSHPQFEK), replaced the antibody sequence. Point mutations to spike protein were subsequently introduced using Quikchange site-directed mutagenesis kit (Agilent). For RBD expression, only amino acids 331-524 were cloned in together with Twin-strep tag and the leader sequence. Purified plasmid was transfected into HEK293 suspension cell culture in F17 media (ThemoFisher) at a concentration of 1.0mg DNA/L of culture with branched polyethylenimine (Sigma-Aldrich) at 3:1 w/w ratio of PEI:DNA. Culture supernatant was harvested after 7 days and purified by HPLC using StrepTactinXT Superflow High Capacity Column (IBA Life GmBH) or MabSelect SuRe Column (Cytiva) for viral proteins or antibodies respectively.

Virus culture and microneutralization

SARS-CoV-2 virus obtained from a patient nasal swab was cultured in VeroE6 cells and supernatant harvested on observation of 90% CPE (hCoV-19/Singapore/3/2020). Antibodies at indicated concentrations was incubated with 100 $TCID_{50}$ of virus and $2x10^4$ VeroE6 cells in 100µl of culture media (MEM/2% FCS) in 96-well flat bottom plates and incubated for 72hrs. Neutralization was measured using Viral Toxglo reagent (Promega) to determine percentage cell survival relative to a no virus and virus only controls. For initial screening of B cell supernatant, 12.5µl of supernatant was mixed with 25 $TCID_{50}$ of virus instead.

### ELISA

To determine antibody affinity to RBD, WT or mutant spike protein, 2ug/ml purified protein in binding buffer (100 mM Tris-HCl, 1 mM EDTA, 150 mM NaCl, pH8.0) was coated onto Streptactin XT 96-well ELISA plates (IBA GmBH) for 2hrs. Plates were washed with PBS and antibody diluted into 2% BSA/PBS blocking solution at indicated concentrations then incubated for 1hr before washing thrice with PBS/0.05%Tween. For binding at different pH, antibodies were incubated in 1xPBS adjusted to the appropriate pH with HCl with 0.5% BSA as block. Antibody binding was detected using 1:5000 anti-huIgG Fc-HRP conjugated secondary antibody (ThermoFisher) diluted in blocking solution incubated for 1hr. Plates were then washed thrice with PBS/0.05% Tween and once with PBS. After washing, plates were developed with colorimetric detection substrate 3,3′,5,5′-tetramethylbenzidine (Turbo-TMB; Pierce). The reaction was stopped with 2M H2SO4, and OD was measured at 450 nm

### Binding inhibition assay

20mM Twin-strep-tagged viral protein was incubated with antibody for 1hr in 50µl FACS buffer. $5x10^4$ huACE2 expressing CHO cells (a kind gift from A/Prof Dr Tan Yee Joo, National University of Singapore) in 50ul FACS buffer was added and incubated for a further 1hr. Cells were spun down at 300g for 5min, washed twice with PBS and then stained with 1:50 Alexa488 conjugated anti-Strep-Tag II antibody (StrepMAb-Immuno, IBA GmBH) for 30min in 50µl FACS buffer. All incubations were carried out at 4°C. Cells were finally washed once with 1x PBS and stained with 1:100 propidium iodine (PI) and analysed on a BD FACS Canto II. PI- cells were gated and binding measured by 488 channel fluorescence intensity.

### Charge Variant Analysis

Briefly, for charge variants, 35 ug protein samples were loaded onto a HPLC system (Thermo Fisher) coupled with an analytical cation exchange column (YMC BioPro SP-F, 4.6 x 100 mm, 5 um) and UV detector (VWD) at UV length 280 nm under 0.8 mL/min constant flowrate. The proteins samples are eluted by pH gradient (25% - 45% Mobile phase B over 22 mins, Mobile phase A: CX-1 pH 5.6, Mobile phase B: CX-1 pH 10.2, Thermo Fisher), with the acidic surface charged protein elutes earlier than basic surface charged molecule. The relative abundance of acidic variants, basic variants and main isoform are reported. Sample are injected in duplicates.

Isoelectric point determination

Isoelectric point was determine using the PA800 Plus system coupled with UV detector (Beckman Coulter), the protein samples were desalted by buffer exchange to 20 mM Tris pH 8.0 using protein concentrator (Amicon, Merck) and mixed with a mixture of 40 mM arginine, 1.6 mM iminodiacetic, 2.4 M urea, and 4.8% pharmalyte 3-10, as well as 0.8% pI marker 10.0, 9.5 and 4.1, at 0.2 mg/mL final concentration. The mixture was then injected into a neutral-coated capillary with one end submerged in anolyte (phosphoric acid) and the other submerged with catholyte (sodium hydroxide). The molecule migrates to its isoelectric point during the focusing step (15 mins, 25 kV) and was followed by a 30 mins mobilisation phase at a voltage of 30 kV. The A280 signals were collected to determine the main peak pI.

Antibody Dependent Cellular Cytotoxicity (ADCC)

Antibody Dependent Cellular Cytotoxicity was tested using a Jurkat reporter cell line stably expressing FcγRIIIa and an NFAT response element driving downstream expression of firefly luciferase as per manufacturer's protocol (ADCC reporter assay, Promega). Target cells were generated by transiently transfecting HEK293 suspension culture with the full-length WT-spike construct including the transmembrane domain but lacking the C-terminal 19 amino acids which contains an endoplasmic reticulum (ER)-retention signal that had been found to reduce incorporation into pseudovirus[3]. Cells were harvested after 72hrs and seeded at 25,000 cells/well and at 1:3 ratio with reporter cells and purified IgG incubated at indicated concentrations. Luminescence was measured after 6hrs incubation.

Antibody Dependent Enhancement

Spike-bearing viral pseudoparticles were produced through co-transfection with the above full length WT-spike construct along with the lentiviral plasmids pMDLg/pRRE, pRSV-REV (a kind gift from Dr Wang-Cheng-I, Singapore Immunology Network) and the luciferase reporter plasmid pHIV-Luc[4] into HEK293 adherent cells and harvested after 4 days. 5µl of pseudovirus-bearing supernatant was mixed with antibody at indicated concentrations and Raji, THP-1 or ACE2 expressing CHO-cells at 25,000cells/well and incubated at 37°C in a $CO_2$ incubator. Media was changed after 24hrs and luminescence expression measured after a further 24hrs by washing the cells in PBS and adding reagent (Luciferase Assay System, Promega)

Efficacy testing for SARS-CoV-2 infection in K18-ACE2 mice

All animal work was monitored by and performed in accordance with the protocol approved by the DSO Institutional Animal Care and Use Committee (IACUC) and the Institutional Biosafety Committee (IBC). B6.Cg-Tg (K18-ACE2)2Prlmn/J) were obtained from Jackson Laboratory (JAX). Mice for the studies were female between 7 and 12 weeks old. To determine the day of peak viral load, following acclimatization of the mice to the isocages, groups of 3 mice were anesthetized individually with 3% isoflurane using the precision vaporizer and infected intranasally (I.N.) with 50µl of $1.2 \times 10^4$ TCID50 of SARS-

CoV-2. Following infection, the mice were transferred to new isocages. On the indicated days three mice were euthanized by carbon dioxide asphyxiation, the lungs harvested, weighed and made to 10%w/v with viral grow medium then mashed through a disposable mesh using a plunger and aliquoted into screw cap tube and stored at -80$^{o}$ C, for later determination of lung viral load by qualitative real-time PCR (qRT-PCR) and cell culture to determine the tissue culture infective dose (TCID), and cytokine/chemokine mRNA expression. To assess therapeutic efficacy, at the indicated time-point mice were anesthetized with 3% isoflurane using a precision vaporizer and treated with indicated concentration of antibody in 200ul PBS by intra-peritoneal (I.P.) injection, and mice were returned to the isocages for recovery. Lungs were harvested for viral load and cytokine/chemokine mRNA expression at the peak virus day. For the survival groups mice were weighed when indicated and returned to their isocage. Any mice that showed >20% weight loss or significant inactivity were euthanized humanly using carbon dioxide asphyxiation.

Determination of lung viral load in infected mice

Lung viral load was determined using Tissue Culture Infection Dose ($TCID_{50}$) in VERO E6 cells, or by real-time Polymerase Chain reaction (RT-PCR) detecting viral RNA (genome copy number; GCN). Briefly for $TCID_{50}$, serially-diluted lung homogenates were incubated with 2x10$^{4}$ Vero E6 cells in total of 100ul of culture media (MEM/2% FCS) in 96-well flat bottom plates and incubated for 5 days. Virus titre, reciprocal to cell viability, was measured using Viral Toxglo reagent (Promega) to determine cell viability relative to uninfected (cells only) controls. $TCID_{50}$ was subsequently determined using Reed-Muench method. To determine the viral GCN, RNA was extracted from lung homogenates using QIAamp Viral RNA mini kit (Qiagen). Detection of viral RNA was achieved using primers and probes targeted against ORF1ab as described in [5] with 7500 Fast Real-Time PCR system (Applied Biosystem). GCN was determined against standard controls included within each RT-PCR run.

Cytokine and chemokine mRNA measurements

RNA was extracted from lung homogenates harvested at 3 dpi using QIAamp Viral RNA mini kit. cDNA was synthesized using High-Capacity cDNA reverse transcription kit (ThermoFisher) with addition of RNase inhibitor (RNaseOUT, ThermoFisher). Cytokine and chemokine expression was determined using TaqMan Fast Universal PCR mastermix (ThermoFisher) with PrimeTime® Standard qPCR assays (Integrated DNA Technologies) for CCL2 (Mm.PT.58.42151692), CXCL10 (Mm.PT.58.43575827), IL1b (Mm.PT.58.41616450), IL6 (Mm.PT.58.10005566), TNF (Mm.PT.58.12575861), IFNγ1 (Mm.PT.58.30132453.g) and normalised to GADPH (Mm.PT.39a.1) levels. Fold change was determined using the $2^{-\delta\delta Ct}$ method comparing anti-SARS-CoV2 specific or isotype control monoclonal antibodies-treated / irrelevant isotype treated mice, to uninfected mice controls.

Cytokine and chemokine protein measurements

Cytokine and chemokine protein levels in mouse serum were determined by ELISA using paired antibodies for mouse IFNγ (Invitrogen, Cat# 88-7314), IL2 (Invitrogen, Cat# 88-

7024), IL6 (Invitrogen, Cat# 88-7064) and CCL2 (R&D Systems, Cat# DY479-05), according to the manufacturer's instructions. Briefly, 384 well plates were coated with 1X capture antibody for 16hr at 4°C. After blocking for 1hr with blocking buffer provided in the kits, mouse serum was added to the plate and incubated for 2hr at room temperature. Plates were washed thrice with washing buffer and incubated for 1hr with detection antibody, followed by 1hr incubation with Streptavidin-HRP. After washing, plates were developed with colorimetric detection substrate 3,3′,5,5′-tetramethylbenzidine (Turbo-TMB; Pierce). The reaction was stopped with 2M H2SO4, and OD was measured at 450 nm.

Efficacy testing in Golden Syrian hamsters

All work with hamsters was performed at the University of Texas Medical Branch (UTMB) in an AALAC-accredited facility and was approved by the UTMB IACUC and IBC. Biosafety Level 3 tasks were performed by appropriately trained personnel in the Galveston National Laboratory. Golden Syrian hamsters (7-8 weeks old, equal sex) were obtained from Envigo. Animals were group housed (three per cage) in micro-isolator cages for the duration of the study. Food and bedding were routinely supplied, changed, and monitored. Prior to study, hamsters were implanted subcutaneously with IPTT-300 programmable transponders for identification and temperature monitoring. On Day 0, following sedation via isoflurane inhalation (1-5%), hamsters were administered 5 x $10^5$ $TCID_{50}$ SARS-CoV-2 (USA_WA1/2020) via intranasal instillation (50 µl per nare). The virus suspension was prepared on the day of challenge from frozen seed stock initially generated (one passage) in Vero E6 cells from lyophilized material provided by the World Reference Center for Emerging Viruses and Arboviruses at UTMB (TVP 23156). Next generation sequencing confirmed 100% consensus sequence-level match to the original patient specimen (GenBank accession MN985325.1). Four hours post-infection, hamsters were administered sterile saline (vehicle control, n=6) or SC31 (n=6) in sterile saline (20 mg/kg) via intraperitoneal injection. Animals were monitored daily for clinical signs of disease including alterations in body weight and subcutaneous body temperature. Animals were scored based on general appearance, activity, and weight loss. Hamsters presenting with severe disease and/or 20% weight loss were humanely euthanized. Nasal cavity samples, collected daily using 0.5 mm Microbrush® Applicators, were placed into 0.1 mL sterile phosphate buffered saline for viral load analysis. Blood was collected on Days -1, 2, and 7 for hematology analysis using the Abaxis VetScan HM5® Hematology Analyzer per manufacturer instructions. At scheduled study termination (or at humane euthanasia, as applicable), the left lung lobe from each animal was collected, homogenized, and clarified via centrifugation. Clarified homogenate was processed for viral load.

Viral load determination during Hamster studies

Viral load was determined in nasal samples and clarified lung homogenate via $TCID_{50}$ and qRT-PCR assays. For $TCID_{50}$ analysis, samples were serially diluted and incubated with 2 x $10^4$ Vero E6 cells in 100 µl of culture medium (MEM/2% FBS) in 96-well flat bottom plates (4 replicate wells per dilution). Each plate contained negative and positive control wells inoculated with culture medium and diluted virus stock, respectively. Cultures were incubated at 37°C/5% $CO_2$ for 96h after which cytopathic effect was measured via microscopic observation. The $TCID_{50}$ for each sample was calculated using the Reed-Muench method. For qRT-PCR analysis, samples (50 μL) were added to TRIzol® LS Reagent (250 μL) and allowed to incubate under ambient conditions for 10 min. Samples were processed to RNA using Zymo Direct-zol™ RNA Mini Prep kits per manufacturer instructions. RNA samples were analyzed via qRT-PCR targeting the SARS-CoV-2 E gene as initially described by the WHO. Real-time analysis was performed using the Bio-Rad CFX96™ Real-Time PCR Detection System. For quantification purposes, viral RNA extracted from the virus seed stock was used.

Histopathology

Remaining lung tissue following removal of sample for viral load analysis was placed in 10% neutral buffered formalin for 24h after which formalin was replaced and samples were allowed to fix for an additional 21 days minimum. Fixed tissues were processed to hematoxylin and eosin-stained slides and examined by a board-certified pathologist at Experimental Pathology Laboratories, Inc. (EPL®) in Sterling, Virginia. Findings were graded from minimal to severe based on a standardized scoring scale.

### Fig. S1

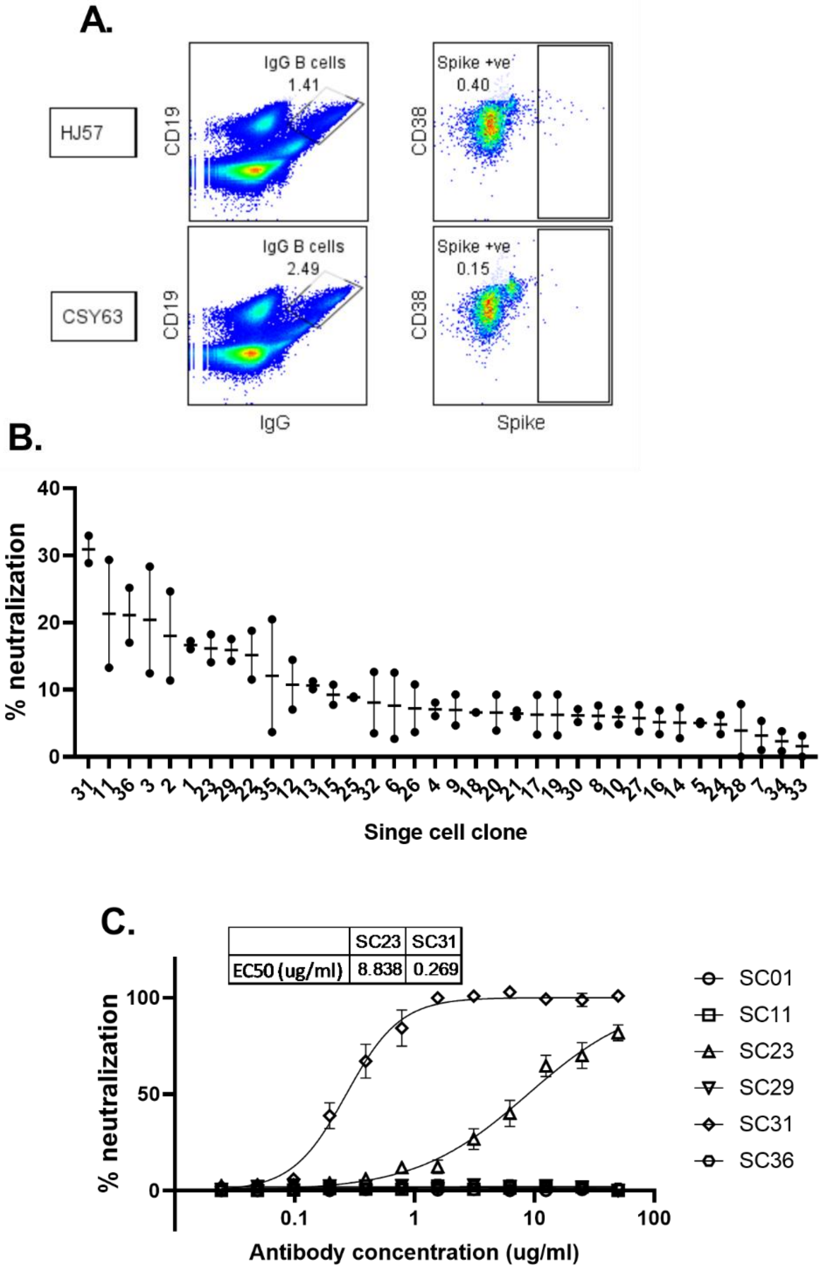


**Fig S1A. Discovery of SC31 from single B cells.** (**A**) Dot plot showing gating strategy for sorting spike-binding IgG B cells from patient PBMCs samples HJ57 and CSY63. (**B**) Neutralization of 25TCID$_{50}$ of live SARS-CoV-2 coronavirus by single B cell culture supernatants showing spike-binding activity. Each supernatant was tested in duplicate and is shown individually. (C) Neutralization efficacy of single cell IgG1 antibodies against 100TCID$_{50}$ live SARS-CoV-2 coronavirus represented as a percentage respective to uninfected and virus infected cell controls. EC50 values for antibodies showing significant neutralization are shown and results represent the mean of four independent replicates with bars indicating standard error.

### Fig. S2

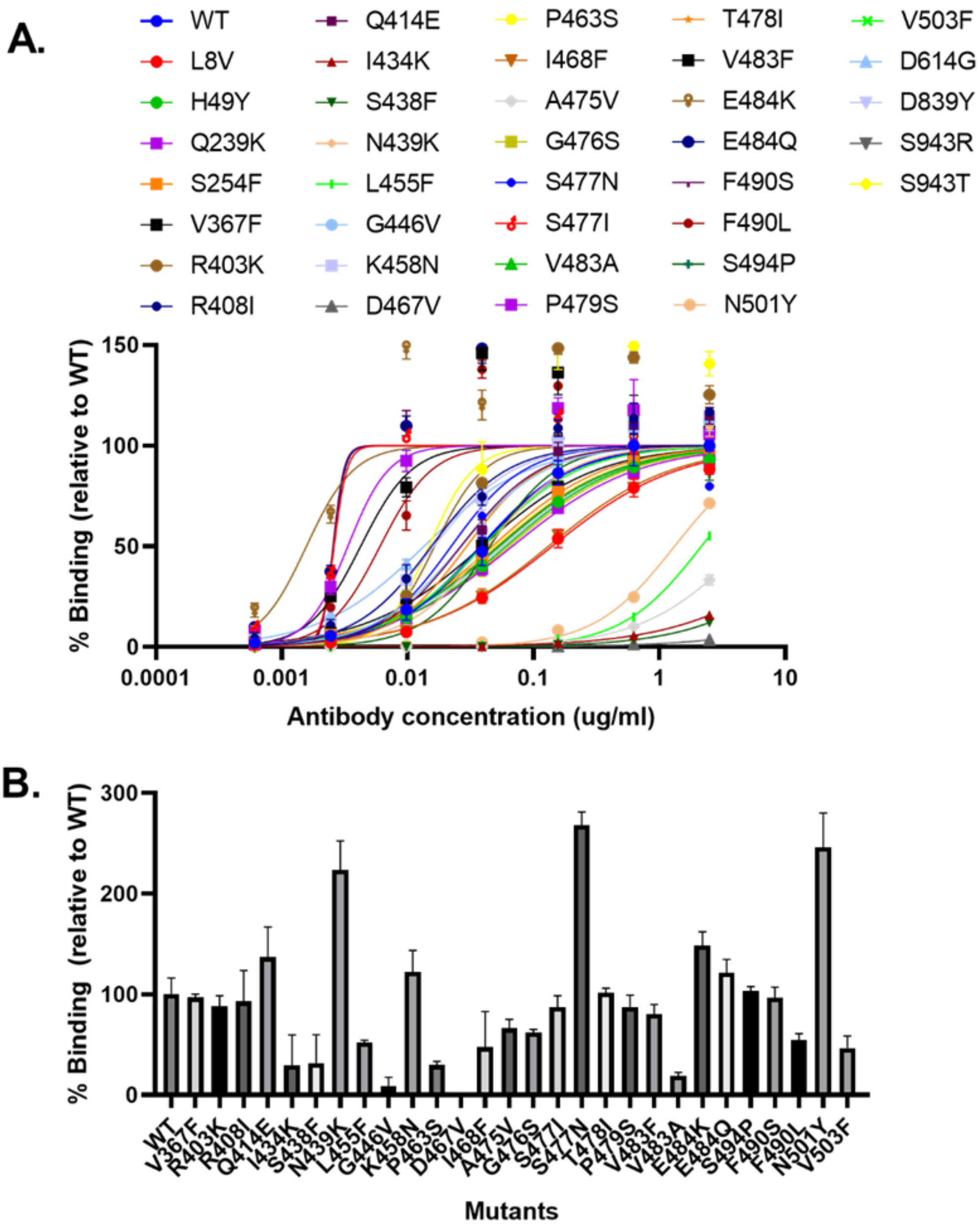


**Fig. S3 Determination of SC31 binding to Spike variants. (A)** Binding affinity of SC31 to purified wild-type spike and spike mutants as determined by ELISA. Results are the mean of three independent replicates and are represented as a percentage of maximal absorbance against wild-type spike at the highest antibody concentration. (**B**) Binding affinity of purified wild-type and mutant spike protein to hACE2-expressing CHO cells as determined by fluorescence intensity with flow cytometry. Results are the mean of three independent replicates with bars showing the standard error and are represented relative to wild-type spike binding to ACE2. Only mutations within the RBD region were tested.

### Fig. S3

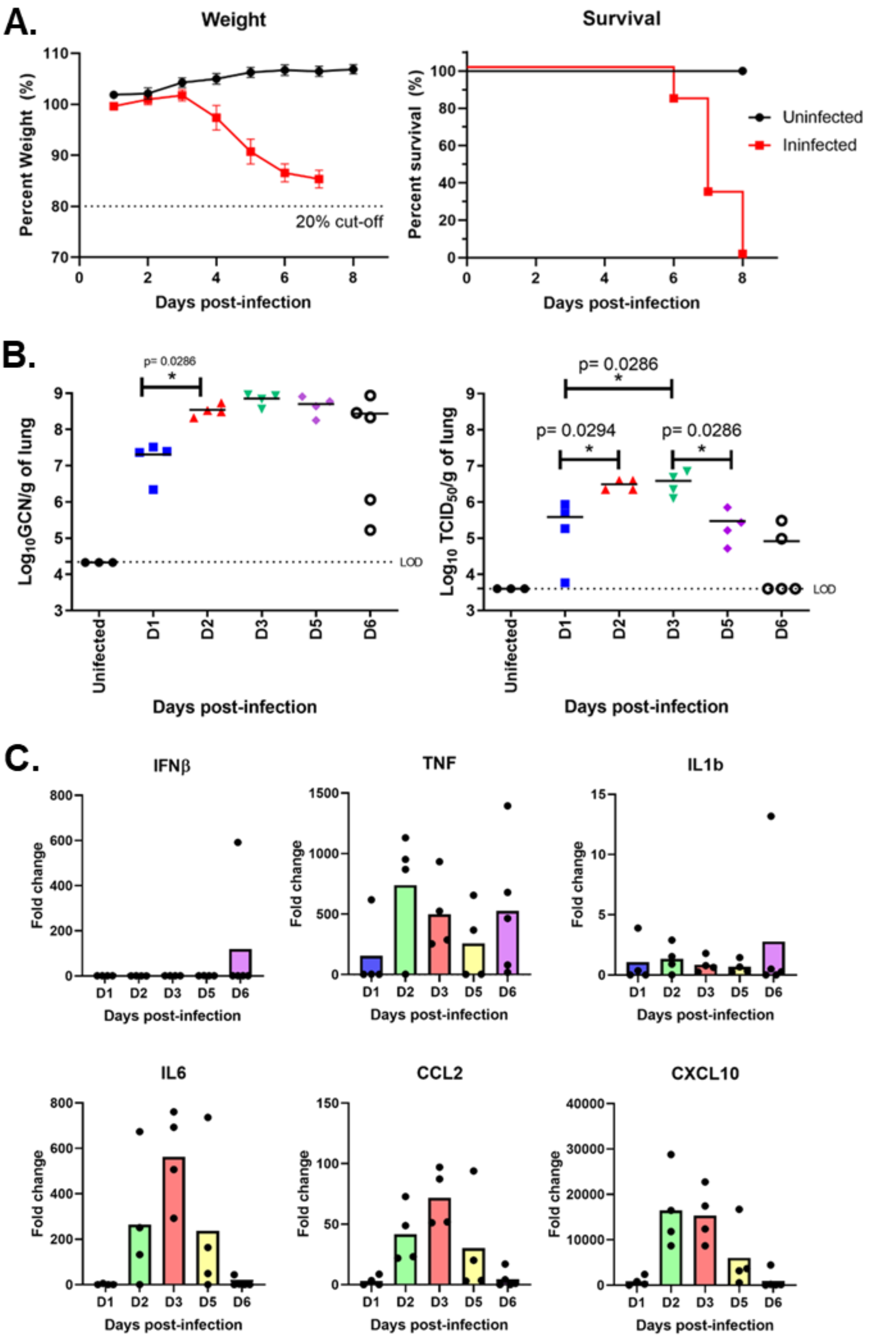


**Fig. S4 Establishment of K-18 human ACE2 transgenic mouse SARS-CoV-2 infection model.** (**A**) Disease progression in K18 mice as shown by weight loss (left) and survival (right). (**B**) Kinetics of viral infection in K18 mice with lung viral load based on genome copies (left) and infectious disease (right). The dotted line indicates the limit of detection (LOD). (**C**). Kinetics of the cytokine response in the lung as measured by mRNA expression of pro-inflammatory cytokines IFNβ, TNF, IL1b, IL6 and chemokines CCL2, CXCL10 represented as fold-change over uninfected mice. Each point represents one individual mouse with the mean indicated by the horizontal lines or bars. Statistical significance between viral load on adjacent days was determined using Student's t-test.

Table S1

**Table S1**: Charge Variant Analysis (CEX-HPLC) and Isoelectric Point (cIEF) of SC31 parental and SC31 engineered demonstrating the improvement in developability

| | **Total Acidic Peaks (%)** | **Main Isoform (%)** | **Total Basic Peaks (%)** | **pI** |
|---|---|---|---|---|
| SC31 parental | 55.6 ± 0.051 | 39.1 ± 0.01 | 5.2 ± 0.03 | 7.9 |
| SC31 engineered | 37.8 ± 0.06 | 57 ± 0.01 | 5.2 ± 0.05 | 8.1 |
